# Molecular mechanisms of PDCD4-mediated modulation of translation initiation and termination

**DOI:** 10.64898/2026.04.14.718391

**Authors:** Walaa Al Sheikh, Ekaterina Shuvalova, Nikita Biziaev, Ali Salman, Petr Kolosov, Alexey Shuvalov, Elena Alkalaeva

## Abstract

PDCD4 is a tumor suppressor, known to affect protein translation by binding to a component of the eIF4F complex, eIF4A, and reducing its helicase activity, which is necessary for the 48S preinitiation complex formation and scanning of the 5’ untranslated region of mRNA. PDCD4 has also been shown to interact with the ribosome and with translation initiation factors eIF4G, eIF4G2, eIF3, and PABP, all of which participate both in initiation and the closed-loop structure that couples initiation and termination. To investigate whether PDCD4 modulates initiation and termination through these interactions, we used a reconstituted mammalian translation system and pre-termination complexes purified from rabbit reticulocyte lysate. We found that PDCD4 suppresses early initiation events prior to eIF4F complex binding to the cap structure on mRNA. Moreover, inhibition of the helicase activity of eIF4A by PDCD4 is lost when the 40S subunit is present. Inhibition of 48S complex formation was also observed in the presence of the truncated eIF4G fragment p50 or the eIF4G2 isoform, both of which interact with eIF4A but lack the eIF4E-binding domain. PDCD4-mediated inhibition of initiation persisted regardless of the presence of PABP. During translation termination, PDCD4 did not affect eIF4A activity, indicating that its regulatory function toward eIF4A is stage-specific and restricted to initiation. Finally, we discovered that PDCD4 additively stimulates peptide release together with eIF3, eIF4G2, and PABP, but competes with eIF4F. Thus, PDCD4 employs a complex molecular mechanism targeting multiple translation factors to regulate different stages of protein synthesis.

## INTRODUCTION

Protein translation consists of four stages: initiation, elongation, termination and ribosome recycling. Each of these stages is controlled by its own set of associated factors. The eukaryotic initiation factor 4F (eIF4F), which consists of eIF4E, eIF4A, and eIF4G, plays a central role in initiation by promoting formation of the 48S pre-initiation complex (PIC) at the 5’ end of capped mRNA [1,2]. Each subunit of this complex performs a specific function – eIF4E binds the 5’ cap of the mRNA, eIF4A is an ATP-dependent RNA helicase that unwinds secondary structures in the 5’ UTR, and eIF4G acts as a scaffold that bring these factors together [3–6]. eIF4G also interacts with initiation factor eIF3, which is associated with the 40S ribosomal subunit, thereby recruiting mRNA to the ribosome [7]. Another eIF4G isoform, eIF4G2 (also known as DAP5), participates in initiation by promoting leader sequence scanning and translation of upstream open reading frames (uORFs) [8,9]. It interacts with eIF4A and eIF3 but lacks the eIF4E binding domain [10–12]. eIF4F facilitates the formation of the circular structure of mRNA (closed-loop) by joining the 5’ cap to the 3’ poly(A) tail through the interaction of eIF4G with poly(A)-binding protein (PABP), which increases translation efficiency [13–17].

The closed-loop also brings the terminating ribosome into proximity with initiation factors. In eukaryotes, two release factors are involved in translation termination, eRF1 and eRF3. eRF1 in the complex with eRF3 recognizes stop codon in the A site of the ribosome [18] and after GTP hydrolysis, performed by eRF3, it hydrolyzes peptidyl-tRNA [19]. eRF3 interacts with PABP, promoting convergence of eRFs and eIF4F. Stimulation activity of eIF4F subunits was revealed in translation termination [20]. eIF4A has been implicated in assisting with the loading of eRF1 onto the ribosome, whereas eIF4G (and it’s truncated form p50) enhances eRF3 GTPase activity. eIF4G2 also stimulates eRF3 GTPase activity [20]. Additionally, eIF3 promotes eRF1 loading and peptide release [21,22]. Thus, initiation and termination are closely linked via closed-loop, and key initiation factors also regulate the end of protein synthesis.

Programmed cell death protein 4 (PDCD4) serves as an important translation regulator with functions in both initiation and termination [23,24] with tumor-suppressor capabilities [25]. It is a highly conserved phosphoprotein capable of shuttling between the nucleus and cytoplasm [26]. Structurally, the protein contains two MA3 domains in its central and C-terminal regions, in addition to an N-terminal domain that participates in RNA binding (Fig. 1A). The MA3 domains bind to eIF4A, with both domains together exhibiting a stronger interaction compared to either domain alone, suggesting that PDCD4 may simultaneously bind two eIF4A molecules [23,27]. PDCD4 primarily inhibits translation initiation by binding to eIF4A, preventing the formation of eIF4F and suppressing the assembly of the 48S complex. Moreover, PDCD4 reduces the overall efficiency of translation initiation, in particular for mRNA with highly structured 5’ UTRs, probably by inhibiting the helicase activity of eIF4A [23]. In addition to binding eIF4A, PDCD4 can also interact with the middle MIF4G domain of eIF4G [28]; thus, it can potentially influence the activity of this protein. Recently, the interactions of PDCD4 with another initiation translation factors eIF3 and eIF4G2 were revealed [29,30]. These findings point to a more complex, multi-factor regulation of translation initiation by this protein. The cryo-electron microscopy studies have provided structural insights into how PDCD4 regulates translation initiation as the structures of PDCD4 in complex with the 40S subunit, both alone and in combination with initiation factors eIF4A, eIF3, and eIF1. These studies revealed that PDCD4’s C-terminal domain interacts with eIF4A at the mRNA entry site on the 40S ribosomal subunit, while its N-terminal domain inserts into the mRNA channel and decoding site. This positioning suggests that PDCD4 blocks the eIF4F-independent functions of eIF4A during mRNA recruitment and scanning, thereby inhibiting translation initiation [29,31].

**Figure 1.**
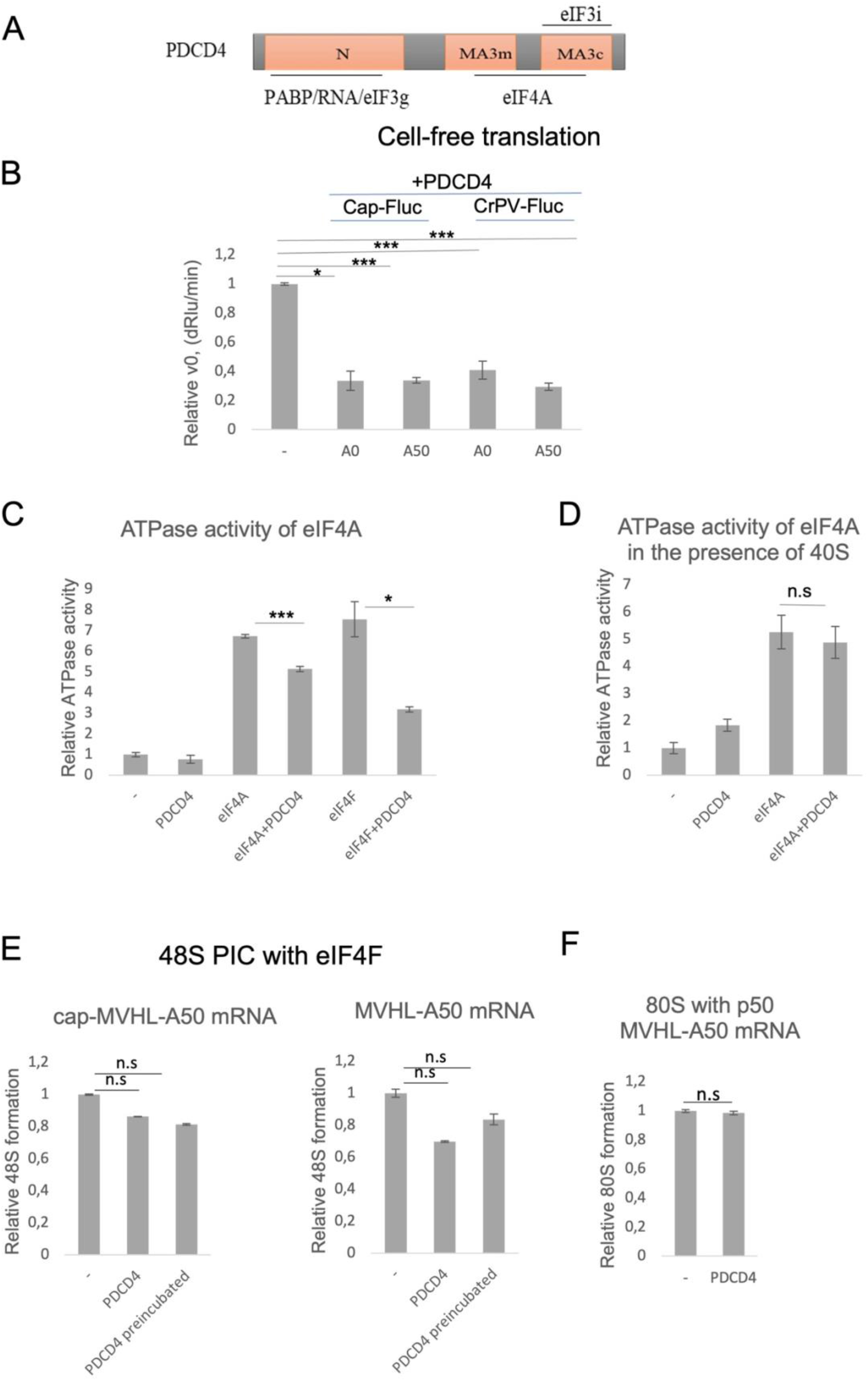
PDCD4 domain structure and its effects on translation and ATPase activity. (A) Domain organization of PDCD4 (RNA-binding domain, MA31, MA32) with the D418A mutation and binding partners indicated. (B) PDCD4 inhibits translation of capped and CrPV IRES-containing Fluc mRNAs in HEK293 lysate. Luminescence normalized to control (no PDCD4). Mean ± SEM (n=3). *, P<0.05; ***, P<0.001. (C) PDCD4 suppresses ATPase activity of free eIF4A and eIF4A within eIF4F. Relative phosphate release. Mean ± SEM (n=3). (D) Addition of 40S ribosomal subunit rescues eIF4A ATPase from PDCD4 inhibition. n.s., not significant; *, P<0.001. (E–F) PDCD4 does not significantly affect 48S PIC formation on capped (E) or uncapped (F) mRNA when eIF4F is present. PDCD4 added directly or preincubated with eIF4F. Normalized 48S signal. Mean ± SEM (n=3). n.s., not significant. (G) PDCD4 does not affect 80S complex formation on uncapped mRNA. Normalized 80S signal. n.s., not significant.

Beyond initiation, PDCD4 also influences termination by modulating the activity of eRF3 [24]. Interacting with the ribosome, it facilitates binding of release factors to the ribosome and dissociation of eRF3 after GTP hydrolysis. This activity is increased in the presence of PABP, as PDCD4 interacts with PABP through RNA-binding motifs present on both proteins [32]. By interfering with eRF3–PABP binding and disrupting new closed-loop formation, PDCD4 may suppress subsequent rounds of translation. Thus, PDCD4 interactions with the ribosome, PABP, and initiation factors (eIF4A, eIF4G, eIF4G2, eIF3) could affect both initiation and termination in the context of the closed-loop.

Because the regulation of initiation and termination by PDCD4 has been described mainly in relation to eIF4A, release factors, and the ribosome, the functional interplay with its other binding partners remains unexplored. Here we used a reconstituted mammalian translation system to dissect the effects of PDCD4 on 48S PIC and 80S ribosome assembly in the presence of eIF4F, eIF4A, eIF4G, p50, eIF4G2, eIF3, and PABP. We also examined the mutual influence of PDCD4 and these factors on translation termination using a peptide release assay. Our results reveal stage-specific regulatory mechanisms of PDCD4.

## MATERIALS AND METHODS

### Purification of ribosomal subunits, translation factors and aminoacylated tRNAs

The 40S and 60S ribosomal subunits and eukaryotic translation factors eIF2 and eIF3 were purified from rabbit reticulocyte and HeLa cell lysates as described previously [33]. Human translation factors eIF1, eIF1A, eIF4A, eIF4B, p50 (MIF4G domain of eIF4G), ΔeIF5B, eIF5, eRF1, eRF1(AGQ) and PABP were produced as recombinant proteins in *E. coli* strain BL21 and purified by Ni-NTA agarose followed by ion-exchange chromatography [33]. Human eRF3a, eIF4F, eIF4G and eIF4G2 were expressed in Sf21 insect cells using the EMBacY baculovirus (MultiBac system) and purified as described [20,34–36]. To obtain eIF4F, all three subunits (eIF4G, eIF4A, and eIF4E) were co-expressed in a baculovirus expression system and the trimer was purified by affinity and ion exchange chromatography. PDCD4 and its D418A mutant were expressed in *E. coli* BL21 cells and isolated as described previously [24].

To obtain aminoacylated total bovine tRNA, a mixture containing 0.25 g/L total bovine tRNA (Sigma, Saint Luis, USA), 0.5 g/L purified ARSs, 100 µM of each amino acid, 40 mM Tris-HCl pH 7.5, 15 mM MgCl_2_, 2 mM DTT, 10 mM ATP, 1 mM CTP, 0.13 u/µL Ribolock (Thermo Fisher Scientific, Waltham, USA) was incubated at 37°C for 7 min. Aminoacylated tRNAs were purified by acidic phenol extraction and gel filtration on NAP-5 columns (Cytiva, Buckinghamshire, UK).

### Model mRNAs

Model mRNAs were synthesized by run-off transcription using the T7 RiboMAX ™ Large Scale RNA Production System kit (Promega, Madison, USA) according to the manufacturer’s protocol. Templates for run-off transcription were generated by PCR amplification of 5’ and 3’ UTRs, and coding sequences from corresponding plasmids (primers listed in Table S1). For mRNAs with a poly(A) tail, the reverse primer contained 50 nt poly(T) sequence was used. To obtain Nluc mRNA, we used pNL-globin plasmid containing UAA stop codon at the end of Nluc coding sequence [37]. To obtain MVHL mRNA, we used pET28-MVHL-UAA plasmid, encoding T7 promoter, four CAA repeats, nt 1–21 of the b-globin 5′ UTR, MVHL tetrapeptide followed by stop codon UAA and 3′ UTR comprising the rest of the natural β-globin coding sequence [38]. To obtain FLUC mRNA containing the IRES sequence of CrPV, the plasmid PGL3R-CPV-FLUC was used [39]. Capped mRNA was synthesized by including 3 mM ARCA (3 mM 3’-O-Me-m^7^G(5’)ppp(5’)G, NEB) at a 3.33-fold excess over GTP. After transcription, mRNA was purified by acidic phenol extraction, precipitated with 3 M LiCl, and washed with 80% ethanol.

### Cell free translation

HEK293F lysate was prepared as described in [40]. 0.25 pmol of IRES FLuc mRNA or cap Fluc mRNA was incubated in 10 μl mixture containing 50% (v/v) HEK293F lysate, 20 mM HEPES–KOH pH 7.5, 2 mM DTT, 0.25 mM spermidine, 0.6 mM Mg (OAc)_2_,16 mM creatine phosphate, 0.06 U/μl creatine kinase (SigmaAldrich), 1 mM ATP, 0.6 mM GTP, 60 mM KOAc, 0.05mM of each proteinogenic amino acid (Promega), 0.2 U/μlRiboLock (Thermo Scientific), 5 mM d-luciferin and 7pmol PDCD4. Luminescence was measured at 30°C using a Tecan Infinite 200Pro (Tecan, Männedorf, Switzerland).

### ATPase assay

ATPase activity of eIF4A was measured with the Malachite Green Phosphate Assay Kit (Sigma, MAK307). Reactions contained 1 µM eIF4A, 1 µM PDCD4, 1 µM eIF4F and 1.5 pmol 40S was added to 75 ng/µL Nluc poly(A) mRNA, and 75 µM ATP in buffer (20 mM Tris-HCl, 15 mM NH_4_Cl_2_, 5 mM MgCl_2_, 5% glycerol, and 1 mM DTT). After 30 min at 37°C, working solution was added and absorbance at 595 nm was measured.

### 48S PIC assembly

To assemble the 48S PIC, 0.375 pmol MVHL mRNA, 1 mM ATP, 0.2 mM GTP, 0.4U/ul Ribolock (Fisher Scientific, Waltham, USA), 1 pmol 40S ribosomal subunit, 3 pmol fmet-tRNA^Met^, and the eukaryotic initiation factors (1 pmol eIF3, 5 pmol eIF2, 2 pmol eIF4B, 2 pmol eIF1, 2 pmol eIF1A) were mixed in the buffer containing (Tris-HCl 20 mM pH 7.5, KAc 50 mM, MgCl_2_ 2.5 mM, DTT 2 mM, spermidine 0.25 mM). Then the mixture was supplemented with one of the following variations: 2 pmol eIF4F, 7 pmol eIF4A and 2 pmol eIF4G, p50 or eIF4G2. 7 pmol PDCD4 /its mutant D418A or PABP was added to the final mixture directly or was preincubated with the eIF4F or its components for 10 min. The final volume was 10 μl. The mixture was incubated for 15 min at 37°C. To assemble 80S complex, 2 pmol of eIF5 and eIF5b, as well as 1 pmol of 60S were added to the 48S PIC. After addition to 7 pmol of PDCD4 the mixture was incubated for 10 min at 37°C.

2 μl buffer, containing 2.5 mM of each dNTPs, 25 mM Tris-HCl pH 7.5, 50 mM KAc, 2 mM DDT, 40 mM MgCl_2_, 250 nM 5’-6-FAM labeled primer 5’-GCATGTGCAGAGGACAGG-3’ (Syntol, Moscow, Russian Federation), 0.625 U AMV reverse transcriptase (NEB, Ipswich, USA), were added to ribosomal complexes. Reverse transcription was performed for 60 min at 37°C. Then FAM-labeled cDNA products were purified by phenol-chloroform extraction. Obtained cDNAs were separated by capillary electrophoresis using standard GeneScan® conditions on an Applied Biosystems® Sanger Sequencing 3500xL Genetic Analyzer (Thermo Fisher Scientific, Waltham, USA).

### Pull-down assay

HisSUMO-tagged PDCD4 or His-tagged initiation factors (eIF4G2, P50,eIF4A)(20 pmol per 1 µL packed Ni-NTA Sepharose resin) was immobilized on resin pre-equilibrated in wash buffer (WB: 25 mM Tris-HCl pH 7.5, 150 mM KCl, 10 mM imidazole) and preincubated for 10 min at room temperature. After washing, 5 pmol of eIF4A, eIF3 or PABP, PDCD4 were added in binding buffer (25 mM Tris-HCl pH 7.5, 150 mM KCl, 2.5 mM MgCl2, 2 mM DTT, 0.25 mM spermidine, 0.2 mM GTP supplemented with MgCl2) and incubated for 30 min at 37°C with agitation. RNase A (1 µL, up to 0.1 g/L final concentration) was added to disrupt any RNA-mediated interactions, followed by incubation for 30 min at room temperature on a rotator (10–20 rpm). The resin was washed three times with WB, then incubated for 1.5 h at room temperature on a rotator with 0.2 pmol Ulp1 protease to elute bound proteins. Eluates were analyzed by Western blot using antibodies against PDCD4 (Bethyl, Texas, USA, a301-107a), eIF4A (Santa Cruz Biotech, Texas, USA, sc-377315), p50/eIF4G1 (CellSignalling, Danvers, USA, 2858S), eIF4G2 (FineTest, Hubei, China FNab02239), eIF3 (Abcam, Cambridge, UK, ab86146) and PABP (abcam, Cambridge, UK, ab21060).

### Termi-Luc assay

Pretermination complex assembly at NLuc mRNA was performed as described previously [41] with minor modifications. A reaction mixture containing nuclease-treated RRL (Green Hectares, Oregon, USA) was supplemented with 30 mM Hepes-KOH pH 7.5, 50 mM KOAc, 1 mM Mg(OAc)_2_, 0.2 mM ATP and GTP, 0.04 mM 20 amino acids (Promega, Madison, USA), 0.5 mM spermidine, 5 ng/μl total rabbit tRNA, 10 mM creatine phosphate, 0.003 u/μl creatine kinase (Sigma-Aldrich, Burlington, USA), 1 mM DTT and 0.2 U/μl Ribolock (Thermo Fisher Scientific, Waltham, MA USA) in a volume of 200 µl. The mixture was preincubated with 1 μM eRF1(AGQ) at 30°C for 10 min, followed by the addition of NLuc mRNA to a final concentration of 8 μg/ml. Next, the mixture was incubated for 1 hour to obtain pretermination complexes (preTCs) with translated Nluc, and after that, KOAc concentration was adjusted to 300 mM, and the mixture was layered on a 10 to 35% linear sucrose gradient in buffer, containing 50 mM HEPES-KOH pH 7.5, 7.5 mM Mg(OAc)_2_, 300 mM KOAc, and 1 mM DTT. The gradient was centrifuged in a SW-55 Ti (Beckman Coulter) rotor at 55000 rpm for 1 h at 4°C. Fractions enriched with preTC were collected by optical density and peptide release activity in the presence of excess amounts of eRFs. PreTC(NL) was aliquoted, and stored at −80 C.

Peptide release was performed in a solution containing 1.5 pM preTC, 50 mM Hepes-KOH pH 7.5, 50 mM KOAc pH 7.0, 0.25 mM spermidine, 1 mM DTT, 0.2 mM GTP, and 1% NLuc substrate (Nano-Glo, Promega, Madison, USA) in the presence of release factors (2.5 nM each eRF1 and eRF3a), 20 nM each PDCD4, p50, eIF4F, and eIF4A, 5 nM of eIF3 and PABP, and 10nM of eIF4G2,. Luminescence was measured at 30°C using a Tecan Infinite 200Pro (Tecan, Männedorf, Switzerland). The peptide release kinetic curves were generated, and the standard deviation was calculated using Microsoft Excel.

### Statistical analysis

All experiments were performed in at least three technical repeats. Data are presented as mean ± standard error (SEM). A two-tailed t-test was used to compare mean values between two groups. The Holm– Bonferroni method was used to counteract the problem of multiple comparisons.

## RESULTS

### PDCD4 suppresses early, but not late, stages of translation initiation

We first tested the activity of recombinant PDCD4 in translation suppression using HEK293 cells lysate on capped or containing Cricket Paralysis Virus (CrPV) IRES Fluc mRNAs with or without poly(A) tail (Fig. 1B). It should be noted that CrPV IRES is capable to recruit directly the ribosome and initiate translation without initiation factors [42]. PDCD4 significantly decreased translation of capped Fluc mRNA (with or without a poly(A) tail) and of CrPV IRES-containing Fluc mRNA (Fig. 1B, S1). Thus, PDCD4 inhibits translation of both cap-dependent and cap-independent mRNAs, suggesting multiple mechanisms of action.

To confirm that our preparation of PDCD4 affects the helicase activity of eIF4A, we measured ATPase activity. PDCD4 suppressed the ATPase activity of free eIF4A as well as eIF4A within the eIF4F complex (Fig. 1C). Strikingly, addition of the 40S ribosomal subunit to the reaction completely rescued eIF4A from PDCD4-mediated inhibition (Fig. 1D). This indicates that once eIF4A engages with the ribosome, it becomes refractory to PDCD4.

We next reconstituted 48S preinitiation complexes on capped and uncapped polyadenylated mRNA in the presence of intact eIF4F. PDCD4 had only a minor, statistically insignificant effect on 48S assembly, regardless of whether it was preincubated with eIF4F or added directly (Fig. 1E, S2A-B). Similarly, when 48S complexes were allowed to progress to 80S ribosomes on uncapped mRNA, PDCD4 did not alter the efficiency of 80S formation (Fig. 1F, S3D). These results suggest that once the eIF4F complex is fully assembled and bound to the mRNA, PDCD4 can no longer suppress initiation.

### PDCD4 inhibits initiation prior to eIF4F-cap interaction and independently of eIF4A binding

To determine whether the protective effect of eIF4F requires the cap-binding subunit eIF4E, we assembled 48S complexes using a mixture of eIF4A and full-length eIF4G in the absence of eIF4E (see domain structures in Fig. 2A). Under these conditions, PDCD4 significantly reduced 48S formation on both capped and uncapped mRNA, with a stronger inhibition on uncapped mRNA (Fig. 2B, S2C-D). Thus, the relief from PDCD4 inhibition observed in the presence of intact eIF4F (Fig. 1E–F) appears to depend on eIF4E and, likely, on stable eIF4F binding to the capped mRNA 5′ end.

**Figure 2.**
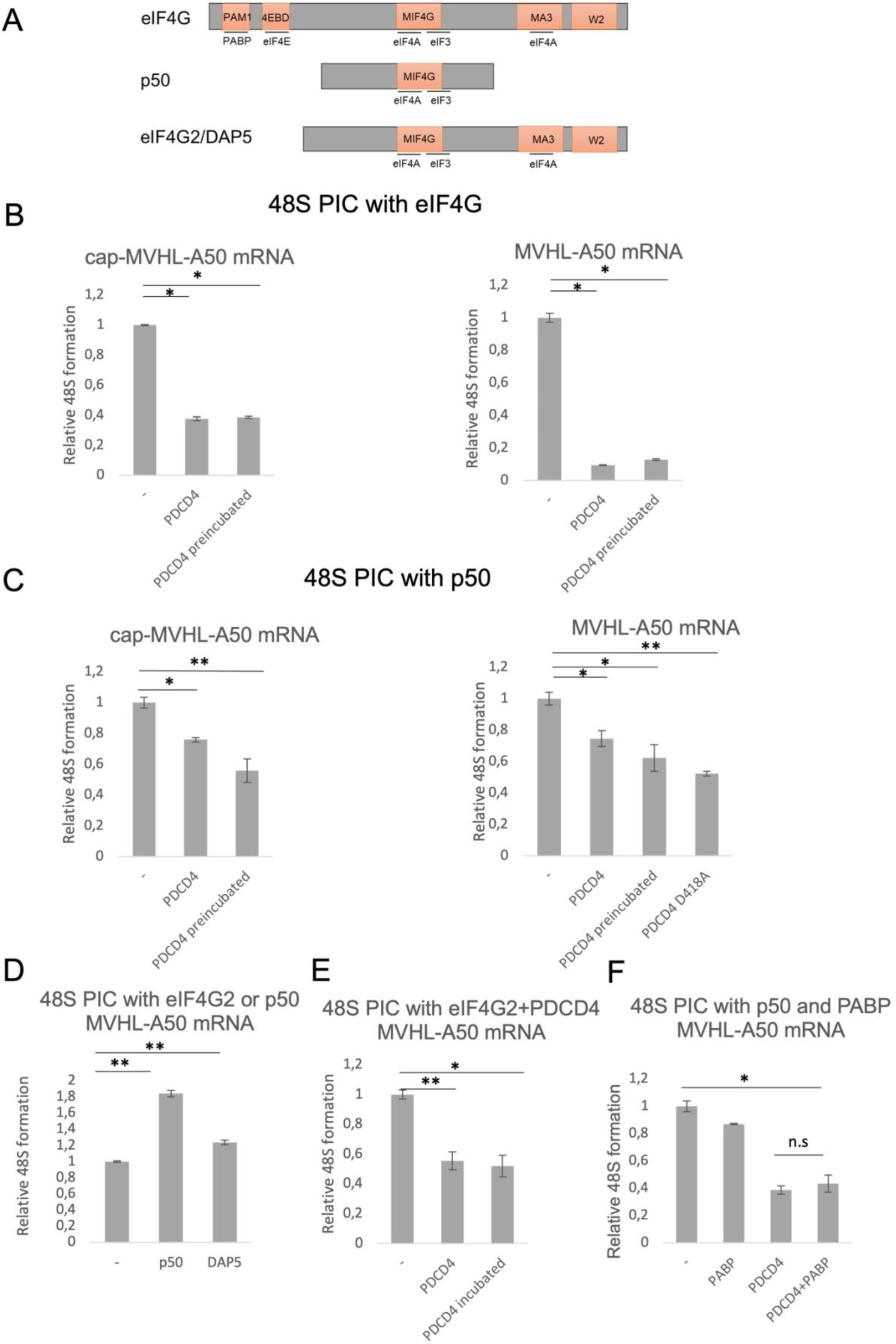
PDCD4 suppresses translation initiation independently of eIF4E and eIF4A binding. (A) Domain schematics of eIF4G, p50 (MIF4G domain), and eIF4G2, showing interactions with eIF4A, eIF4E, PABP, and the ribosome. (B) PDCD4 inhibits 48S PIC assembly when eIF4A and full-length eIF4G (without eIF4E) are used, with stronger effect on uncapped mRNA. Normalized 48S signal. Mean ± SEM (n=3). **, P<0.01; ***, P<0.001. (C) PDCD4 inhibits 48S assembly with eIF4A and p50 on both capped and uncapped mRNA. **, P<0.01. (D) eIF4G2 and p50 stimulate 48S assembly on uncapped mRNA (relative to no eIF4G2). *, P<0.05. (E) PDCD4 inhibits eIF4G2-stimulated 48S formation. **, P<0.01. (F) PABP does not counteract PDCD4 inhibition of 48S assembly on uncapped mRNA. *, P<0.05; n.s., not significant.

We then used p50, a truncated form of eIF4G that binds eIF4A but cannot interact with eIF4E or PABP (Fig. 2A). p50 stimulates 48S assembly on unstructured 5′ UTRs independently of the cap [39]. PDCD4 inhibited 48S complex formation in the presence of p50 and eIF4A by approximately twofold, both on capped and uncapped mRNA (Fig. 2C, S2E-F). Surprisingly, a PDCD4 mutant (D418A) that cannot bind eIF4A also inhibited 48S assembly to a similar extent (Fig. 2C, right panel, S3B). This indicates that PDCD4 possesses an eIF4A-independent inhibitory mechanism, likely involving direct interaction with the ribosome or with p50.

Another eIF4G isoform, eIF4G2, also reported to interact with PDCD4 [30]. In our reconstituted system, eIF4G2 stimulated 48S assembly on uncapped mRNA, albeit more weakly than p50 (Fig. 2D, S2G). PDCD4 suppressed this eIF4G2-dependent stimulation by about half (Fig. 2E, S3C), suggesting that a similar eIF4A-independent mechanism operates with eIF4G2.

Finally, PABP, which interacts with both eIF4G and PDCD4, did not counteract PDCD4 inhibition. In the presence of PABP, PDCD4 still reduced 48S assembly on uncapped polyadenylated mRNA (Fig. 2F, S3A). Thus, PABP does not protect against PDCD4-mediated suppression during initiation.

### PDCD4 directly interacts only with eIF4A among the tested factors

The functional assays suggested that PDCD4 might physically interact with multiple initiation factors. To test this directly, we performed pull-down experiments using immobilized His-SUMO-tagged PDCD4 or His-tagged eIFs. His-SUMO-tagged PDCD4 was incubated with purified eIF4A, eIF3, or PABP, and bound proteins were eluted and detected by Western blotting. His-tagged p50, eIF4G2, and eIF4A were incubated with purified PDCD4 and after elution bound PDCD4 was detected by Western blotting. Under our experimental conditions, PDCD4 specifically pulled down eIF4A, but no binding was detected with p50, eIF4G2, eIF3, or PABP (Fig. 3). Control reactions without PDCD4 or with untagged PDCD4 showed no signals, confirming specificity.

**Figure 3.**
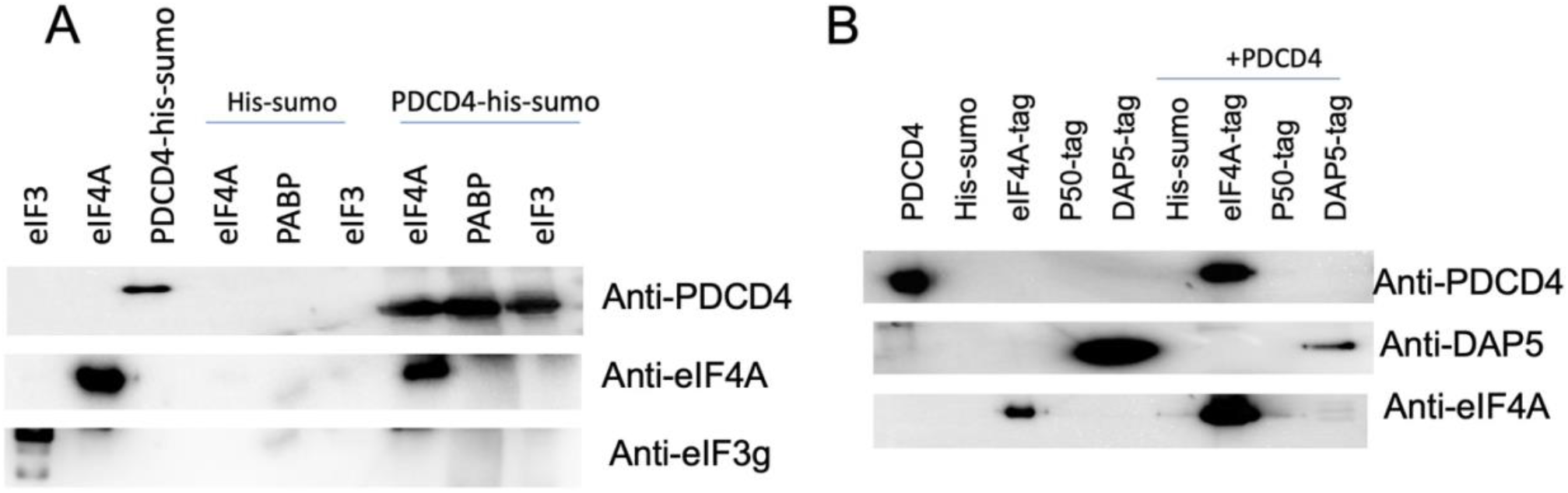
PDCD4 directly interacts only with eIF4A *in vitro*. Pull-down assays with immobilized His-SUMO-PDCD4 or His-tagged factors. Bound proteins were detected by Western blot (antibodies indicated). PDCD4 pulled down eIF4A but not p50, eIF4G2, eIF3, or PABP. Control reactions (no PDCD4 or untagged PDCD4) were negative. Representative of three independent experiments.

These results demonstrate that among the five factors tested, only eIF4A forms a stable direct complex with PDCD4 *in vitro*. The lack of detectable binding to p50 and eIF4G2 suggests that the functional inhibition observed in the presence of these proteins (Fig. 2C, E) is unlikely to be explained by a stable direct PDCD4-p50 or PDCD4-eIF4G2 interaction. Instead, PDCD4 may exert its effects on p50- and eIF4G2-dependent initiation indirectly, for example, through competition for eIF4A binding or through direct interaction with the 40S ribosomal subunit, as suggested by structural studies [23,29]. Similarly, the absence of PDCD4-PABP binding indicates that PABP does not mediate PDCD4 inhibition via a direct contact, consistent with the functional data showing that PABP does not counteract PDCD4 (Fig. 2F).

### PDCD4 differently regulates initiation factors activity during translation termination

Previous studies showed that both PDCD4 and the initiation factors examined here can stimulate activity of eRFs in peptide release [20,21,24,34]. Using the Termi-Luc assay, we asked whether PDCD4 modulates activities of these initiation factors during termination. We measured the initial rate of Nluc release from preTCs in the presence of low concentrations of eRF1/eRF3a (Fig. S4), with or without added factors.

eIF4F alone enhanced peptide release, but when combined with PDCD4, the rate was no higher than with eIF4F alone (Fig. 4A), suggesting that PDCD4 and eIF4F compete for a common target or that eIF4F displaces PDCD4 from the terminating ribosome. In contrast, eIF4A and PDCD4 together stimulated release more than either alone (Fig. 4B), indicating independent action—consistent with eIF4A facilitating eRF1 recruitment and PDCD4 acting at a later step (eRF3 dissociation).

**Figure 4.**
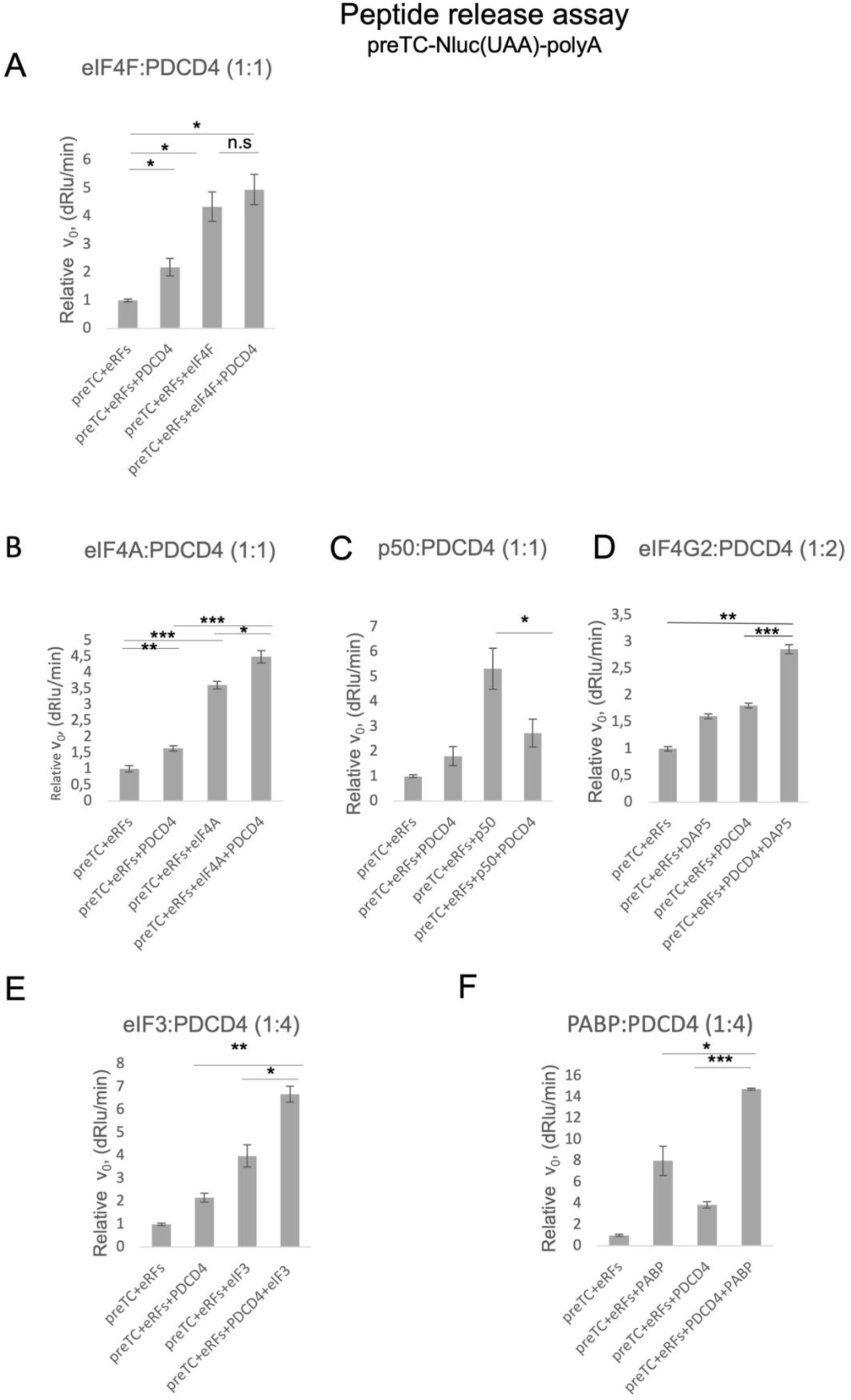
PDCD4 differentially modulates translation termination. Termi-Luc assays on Nluc preTCs (UAA stop codon). Peptide release was induced by eRF1/eRF3a (eRFs) at basal level. Initial rates (v0) normalized to eRFs alone (set to 1). Mean ± SEM (n=3). *, P<0.05; **, P<0.01; ***, P<0.001; n.s., not significant. (A) eIF4F and PDCD4 together do not exceed eIF4F alone. (B) eIF4A and PDCD4 act additively. (C) PDCD4 suppresses p50 stimulation. (D) eIF4G2 and PDCD4 act additively. (E) eIF3 and PDCD4 act additively. (F) PABP and PDCD4 act additively.

p50 also stimulated termination, but adding PDCD4 together with p50 reduced the rate to the level of PDCD4 alone (Fig. 4C). Thus, PDCD4 suppresses p50 activity, likely competing for the same stage (GTP hydrolysis/eRF3 release). Surprisingly, eIF4G2 and PDCD4 together gave an additive effect (Fig. 4D), implying that eIF4G2 may function differently from p50 during termination.

eIF3 and PDCD4 as well as PABP and PABP also acted additively (Fig. 4E-F), consistent with eIF3 and PABP promoting eRF1 loading and PDCD4 acting later. Thus, PDCD4 does not interfere with PABP- and eIF3-mediated enhancement of termination.

Collectively, these results show that during termination PDCD4 acts synergistically with factors involved in stop codon recognition (eIF4A, eIF3, PABP) but competes with eIF4G (p50), which functions at the same step as PDCD4. eIF4G2 appears to behave differently, possibly due to distinct interactions with the ribosome or release factors.

## DISCUSSION

In this study, we dissected the stage-specific functions of PDCD4 in mammalian translation using reconstituted system. Our findings reveal that PDCD4 acts as a checkpoint regulator during early initiation and as a differential modulator during termination, depending on its partners.

PDCD4 inhibited 48S PIC formation when initiation was driven by eIF4A together with eIF4G, p50, or eIF4G2, but had little effect when intact eIF4F was used. This suggests that once eIF4F is fully assembled and bound to the cap, the initiation complex becomes resistant to PDCD4. The ATPase assays support this model: PDCD4 suppressed eIF4A activity in solution, but addition of 40S subunits completely relieved inhibition. Thus, PDCD4 likely targets eIF4A before it engages with the ribosome as part of the scanning machinery. This idea is consistent with cryo-EM structures showing that PDCD4 binds the 40S subunit at the mRNA entry channel, where it contacts a second copy of eIF4A not associated with eIF4G [29,31]. The D418A mutant, which cannot bind eIF4A, still inhibited initiation with p50, indicating an additional eIF4A-independent mechanism—probably direct occlusion of the mRNA channel by PDCD4’s N-terminal domain [31].

A recent study by Dezi et al. reported that PDCD4 can interact with eIF4G2 and eIF3 in cell lysates [30]. The discrepancy with our pull-down results likely arises from differences in experimental conditions, such as the use of purified components versus cell lysates, and suggests that in living cells these interactions may be indirect or mediated by additional factors.

PDCD4 also suppressed translation of CrPV IRES-containing mRNA, which does not require any initiation factors. This observation, together with the ribosome binding reported by others [29,31], argues that PDCD4 can directly interfere with ribosome function, for example by blocking the mRNA entry channel. This may explain its ability to inhibit translation of mRNAs with highly structured 5′ UTRs and certain IRES elements.

During termination, PDCD4 stimulates peptide release by accelerating eRF3 GTP hydrolysis and dissociation [24]. eIF4A, eIF3, and PABP promote the preceding step – eRF1 recruitment and stop codon recognition [20,21,34]. Accordingly, PDCD4 acted additively with these factors. In contrast, p50 (the MIF4G domain of eIF4G) enhances eRF3 GTPase activity [20], the same step targeted by PDCD4; therefore they competed, and PDCD4 suppressed stimulation induced by p50. Interestingly, eIF4G2, which also binds eIF4A and eIF3 but lacks the PABP-interacting region, synergized with PDCD4, suggesting that eIF4G2 may have a different mode of action during termination, possibly involving additional contacts with the ribosome or release factors.

Taken together, our findings reveal that PDCD4 employs distinct mechanisms to regulate initiation and termination, depending on the specific translation factors and the stage of the process. Based on these observations, we propose the following model for PDCD4 action (Fig. 5). During initiation, it surveys eIF4A activity before stable PIC formation, blocking helicase function and mRNA recruitment through both eIF4A-dependent and eIF4A-independent (ribosome-binding) mechanisms. Once eIF4F is assembled and the ribosome is committed, PDCD4 inhibition is lifted. During termination, PDCD4 promotes activity of release factors and dissociation of eRF3, cooperating with factors that stimulate stop codon recognition by eRF1 but competing with those that share its GTPase-activation function. By stimulating termination and preventing reassembly of the closed-loop, PDCD4 may suppress subsequent rounds of translation, contributing to its tumor-suppressive effects.

**Figure 5.**
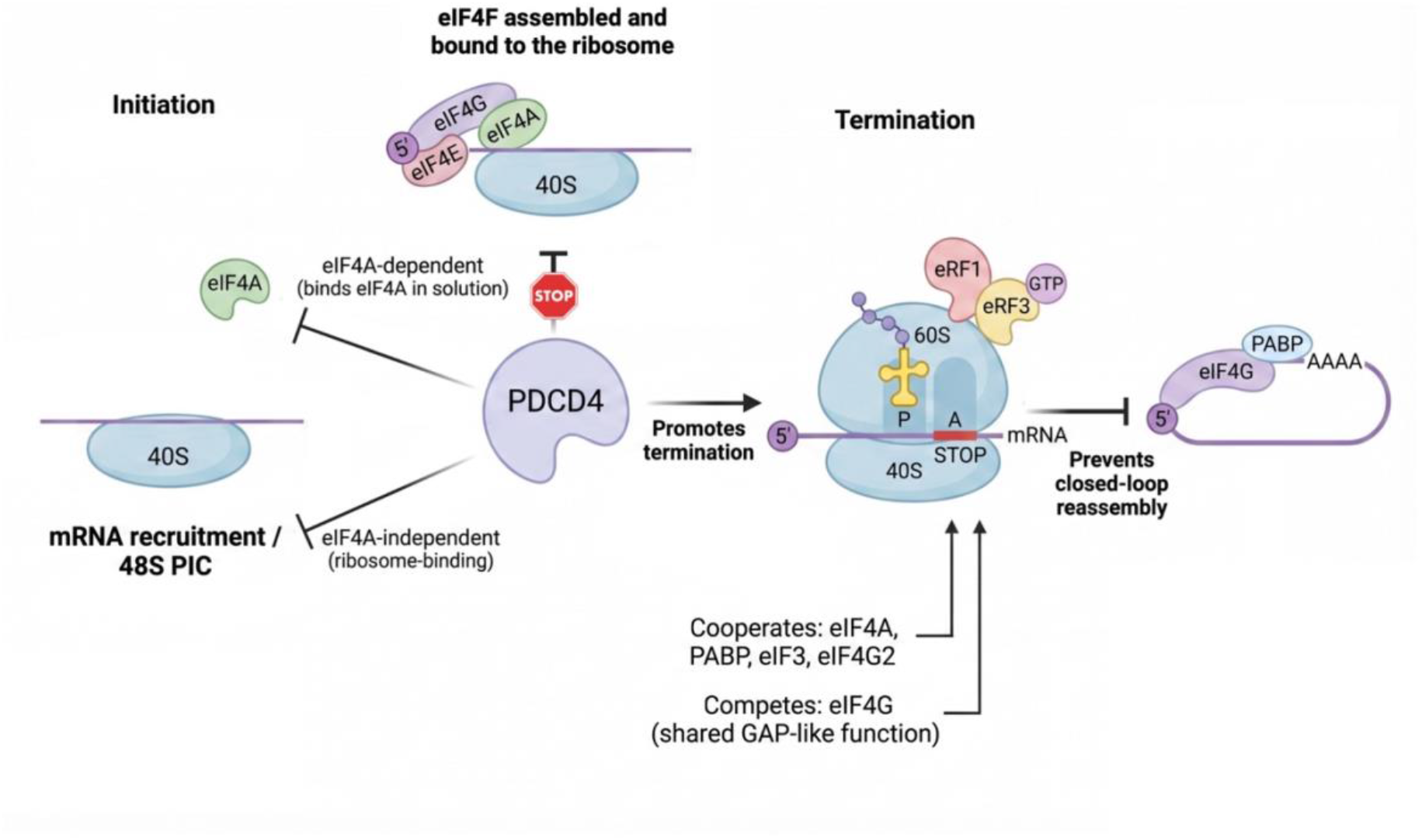
Model of PDCD4-mediated regulation of translation initiation and termination. During early initiation, PDCD4 binds eIF4A in solution (blocking eIF4F formation and helicase activity) and also binds the 40S subunit at the mRNA entry channel (blocking recruitment and scanning). Once eIF4F is assembled and cap-bound, inhibition is lifted. During termination, PDCD4 promotes eRF3 GTP hydrolysis and dissociation, acting additively with factors stimulating loading of eRF1 to the ribosome at the stage of stop codon recognition (eIF4A, eIF3, PABP) but competing with eIF4G (p50). eIF4G2 synergizes with PDCD4.

While this model explains our key observations, certain limitations of our experimental system should be acknowledged. Our *in vitro* system uses short, unstructured leader sequences, which may not fully recapitulate the scanning of long, structured 5′ UTRs or uORF-containing mRNAs where eIF4G2 is particularly important. Future studies should explore PDCD4 effects in more complex cellular contexts and investigate how post-translational modifications (e.g., phosphorylation) modulate its interactions with the ribosome and translation factors. Addressing these questions will provide a more complete understanding of how PDCD4 integrates into the cellular translation regulatory network.

## AUTHOR CONTRIBUTIONS

W.A., E.Sh., N.B., A.S., P.K., and A.Sh. conducted the experiments; E.A. designed the experiments and wrote the paper.

## DECLARATION OF INTERESTS

The authors declare no competing interests.

## SUPPLEMENTAL INFORMATION

Supplementary figures. Figures S1–S4.

Supplementary data. Supplementary figure legends.

## FUNDING

This work was supported by the Russian Science Foundation (Grant No. 22-14-00279)

## ACKNOWLEDGEMENT

We are grateful to Ludmila Frolova for providing us with plasmids encoding release factors, Tatyana Pestova and Christopher Hellen for providing us with plasmids encoding initiation factors. cDNA fragment analyses were performed by the Center of the collective use “Genome” of EIMB RAS.

## Notes

### Competing Interest Statement

The authors have declared no competing interest.

