## Supplemental figures for "Molecular mechanisms of PDCD4-mediated modulation of translation initiation and termination"

#### SUPPLEMENTARY FIGURES LEGENDS

**Figure S1.** The luminescence curves show the effect of PDCD4 (7pmol) on translation of 0.25 pmol of (A) Cap-Fluc-A0 mRNA. (B) Cap-Fluc-A50 mRNA (C) CrPV-Fluc A0 mRNA (D) CrPV-Fluc A50 mRNA in HEK293 cells lysate.

**Figure S2.** Fluorescent toe-printing of 48S complexes, assembled on uncapped and capped mRNA. A representative results of 48S preinitiation complex formation in the reconstituted mammalian translation system on uncapped and capped mRNA given by (A,B) eIF4F (2 pmol), (C,D) eIF4G (2 pmol) and eIF4A (7 pmol), (E,F) p50 (2 pmol) and eIF4A (7 pmol) in the presence/absence of PDCD4 (7 pmol) which was either added directly to reaction or preincubated with the mentioned proteins for 10 min on ice. (G) 48S complexes, assembled on uncapped mRNA, in the presence of p50 or eIF4G2.

**Figure S3.** Fluorescent toe-printing of 48S complexes, assembled on uncapped mRNA. A representative result of 48S preinitiation complex formation in the reconstituted mammalian translation system on uncapped mRNA with p50 (2 pmol) and eIF4A (7 pmol) (A) in the presence/absence of PDCD4/PABP (7 pmol each). (B) in the presence/absence of PDCD4 mutant D418A (7 pmol). (C) 48S complexes, assembled on uncapped mRNA, with eIF4G2 in the presence/absence of PDCD4. (D) A representative results 80S preinitiation complex formation in the reconstituted mammalian translation system on uncapped mRNA with p50 (2 pmol) and eIF4A (7 pmol) in the presence/absence of PDCD4 (7 pmol).

**Figure S4. (**A) Titration curve for release factors. The curve represents the rate of peptide release ($v_0$) at Nluc preTC, induced by eRF1 and eRF3a at different concentrations (2.5 – 20 nM). (B-G) The luminescence curves show the Nluc release induced by eRF1 and eRF3a (2.5 nM each) in the presence of (B) PDCD4 and eIF4F (20 nm each), (C) PDCD4 and eIF4A (20 nm each) (D) PDCD4 and p50 (20 nm each), (E) PDCD4 (20 nm) and eIF4G2 (10 nm), (F) PDCD4 and eIF3 (20 nm PDCD4 and 5nM eIF3), (G) PDCD4 (20 nm) and PABP (5 nm).

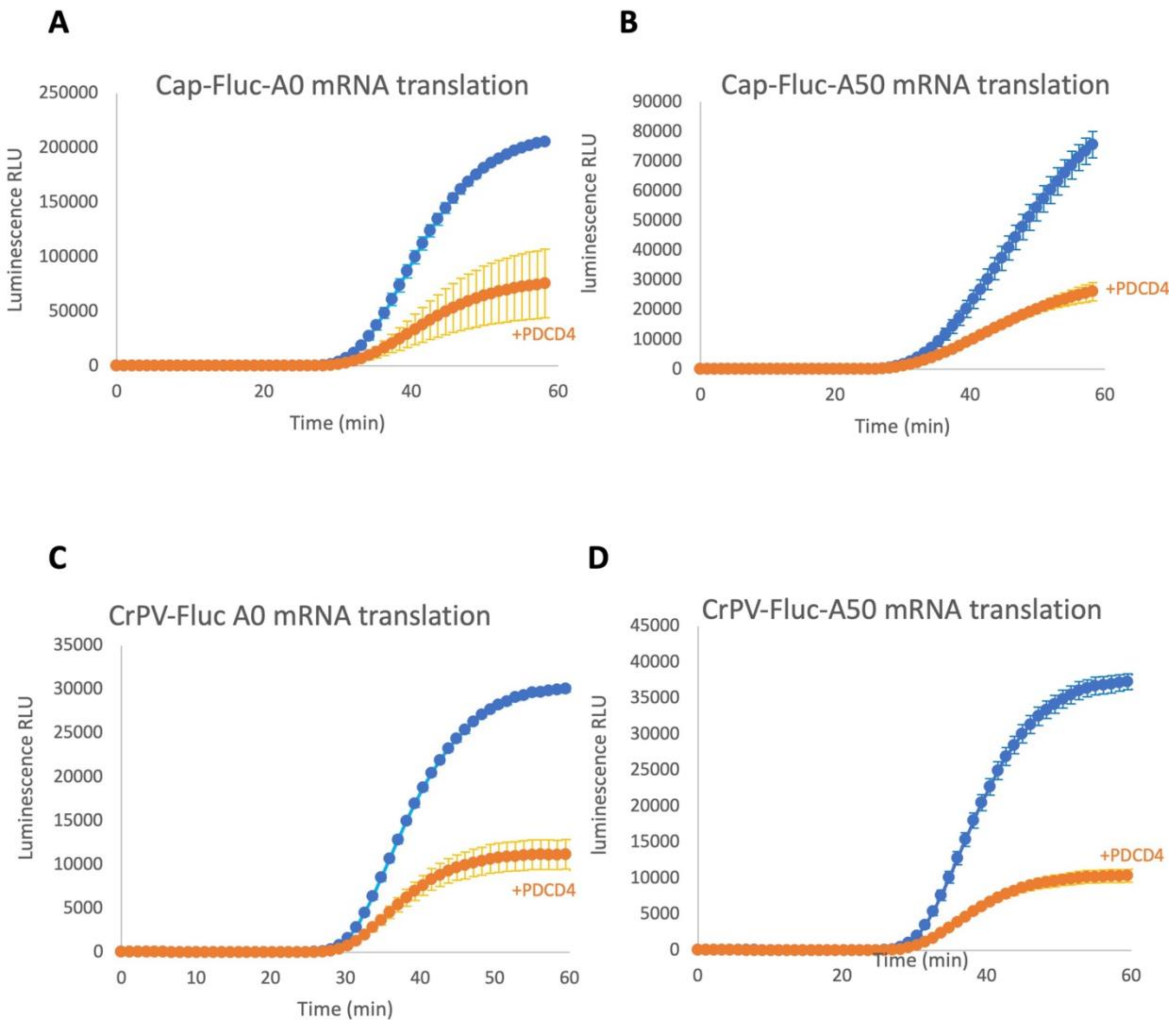
A
Cap-Fluc-A0 mRNA translation
Luminescence RLU
250000
200000
150000
100000
50000
0
0
20
40
60
Time (min)
+PDCD4
B
Cap-Fluc-A50 mRNA translation
luminescence RLU
90000
80000
70000
60000
50000
40000
30000
20000
10000
0
0
20
40
60
Time (min)
+PDCD4
C
CrPV-Fluc A0 mRNA translation
Luminescence RLU
35000
30000
25000
20000
15000
10000
5000
0
0
10
20
30
40
50
60
Time (min)
+PDCD4
D
CrPV-Fluc-A50 mRNA translation
luminescence RLU
45000
40000
35000
30000
25000
20000
15000
10000
5000
0
0
20
40
60
Time (min)
+PDCD4

Fig.S1

### 48S assembly efficiency

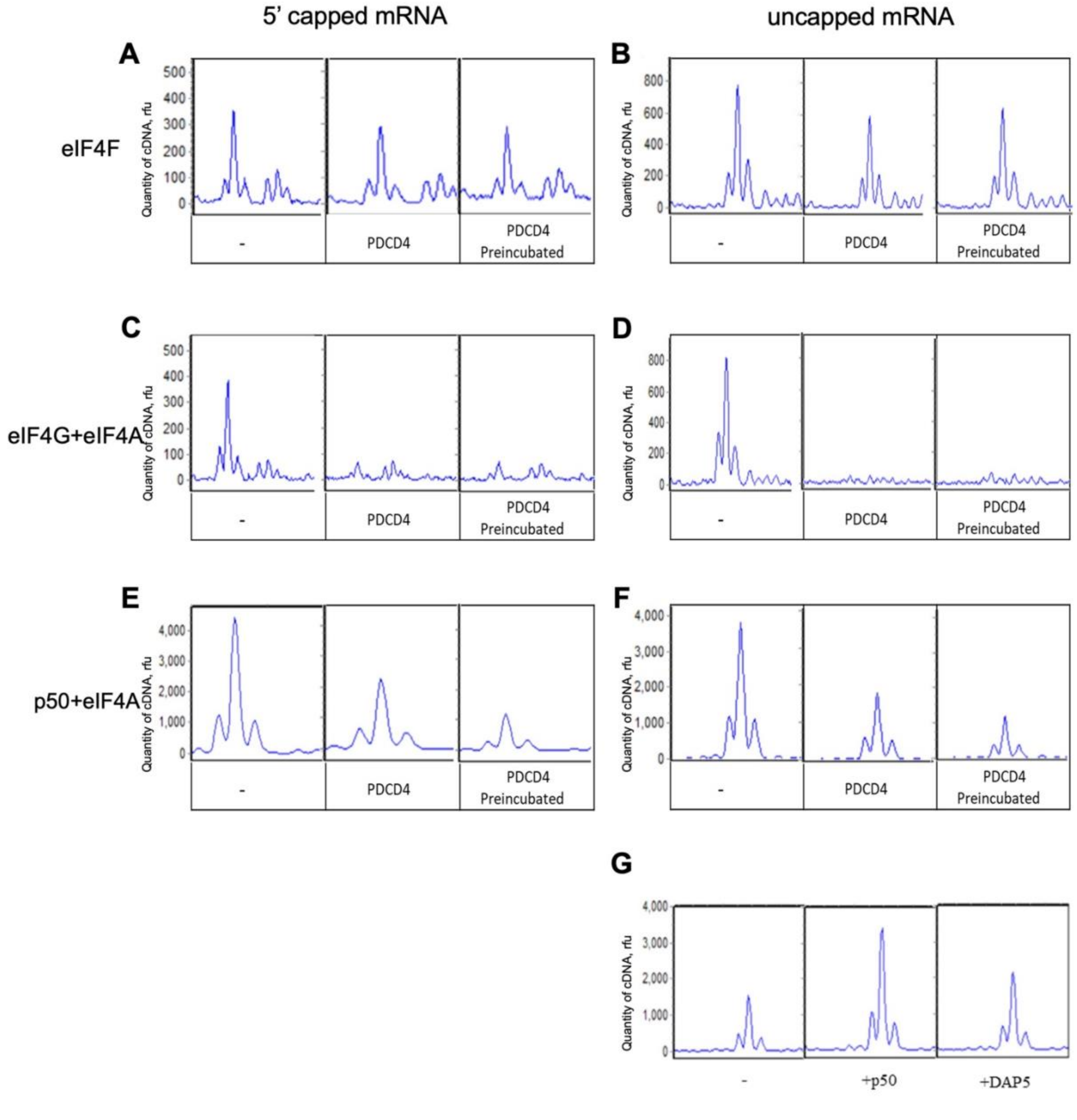


Fig. S2

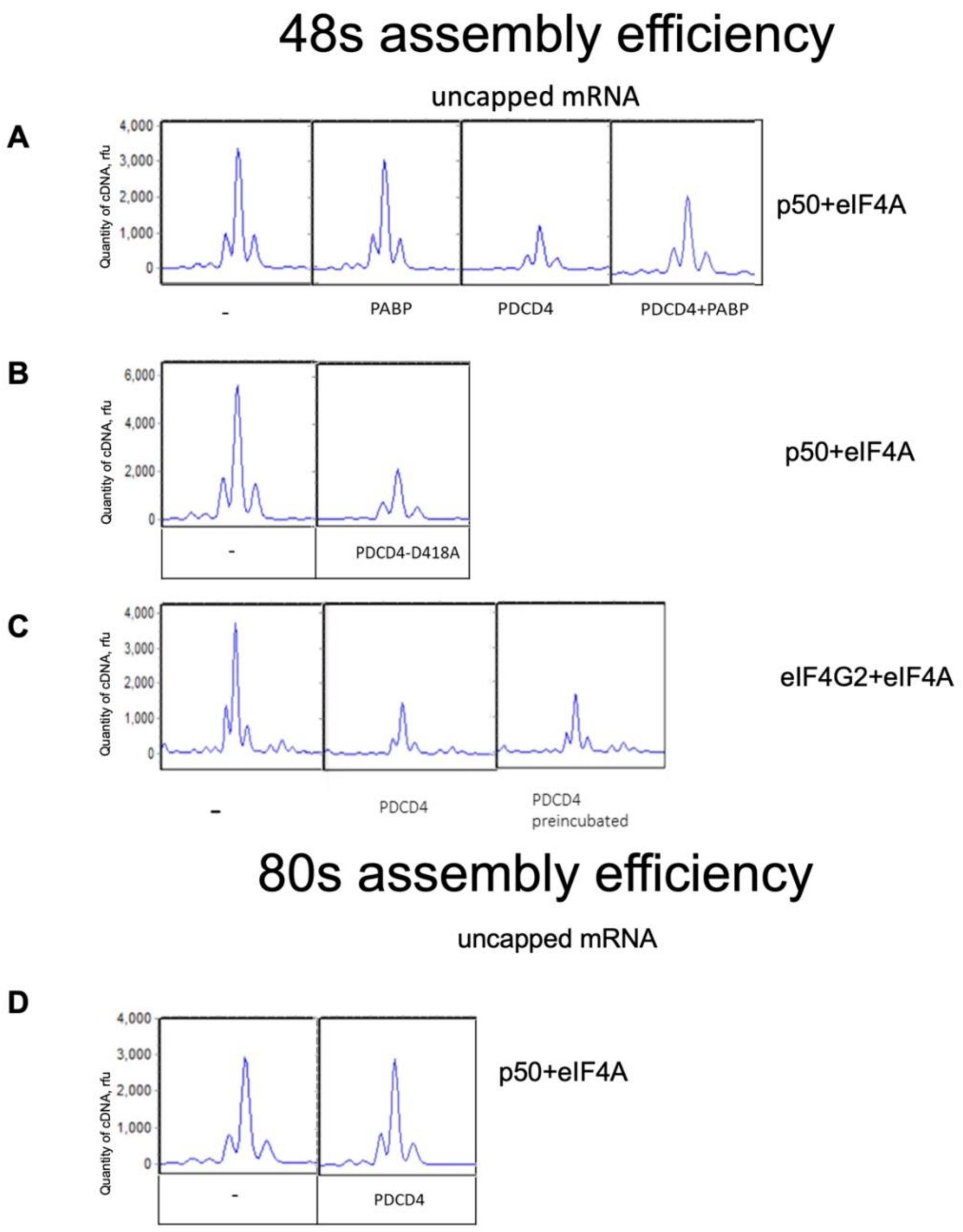
48s assembly efficiency
uncapped mRNA
A
Quantity of cDNA, rfu
4,000
3,000
2,000
1,000
0
-
PABP
PDCD4
PDCD4+PABP
p50+eIF4A
B
6,000
4,000
2,000
0
-
PDCD4-D418A
p50+eIF4A
C
4,000
3,000
2,000
1,000
0
-
PDCD4
PDCD4 preincubated
eIF4G2+eIF4A
80s assembly efficiency
uncapped mRNA
D
4,000
3,000
2,000
1,000
0
-
PDCD4
p50+eIF4A

Fig. S3

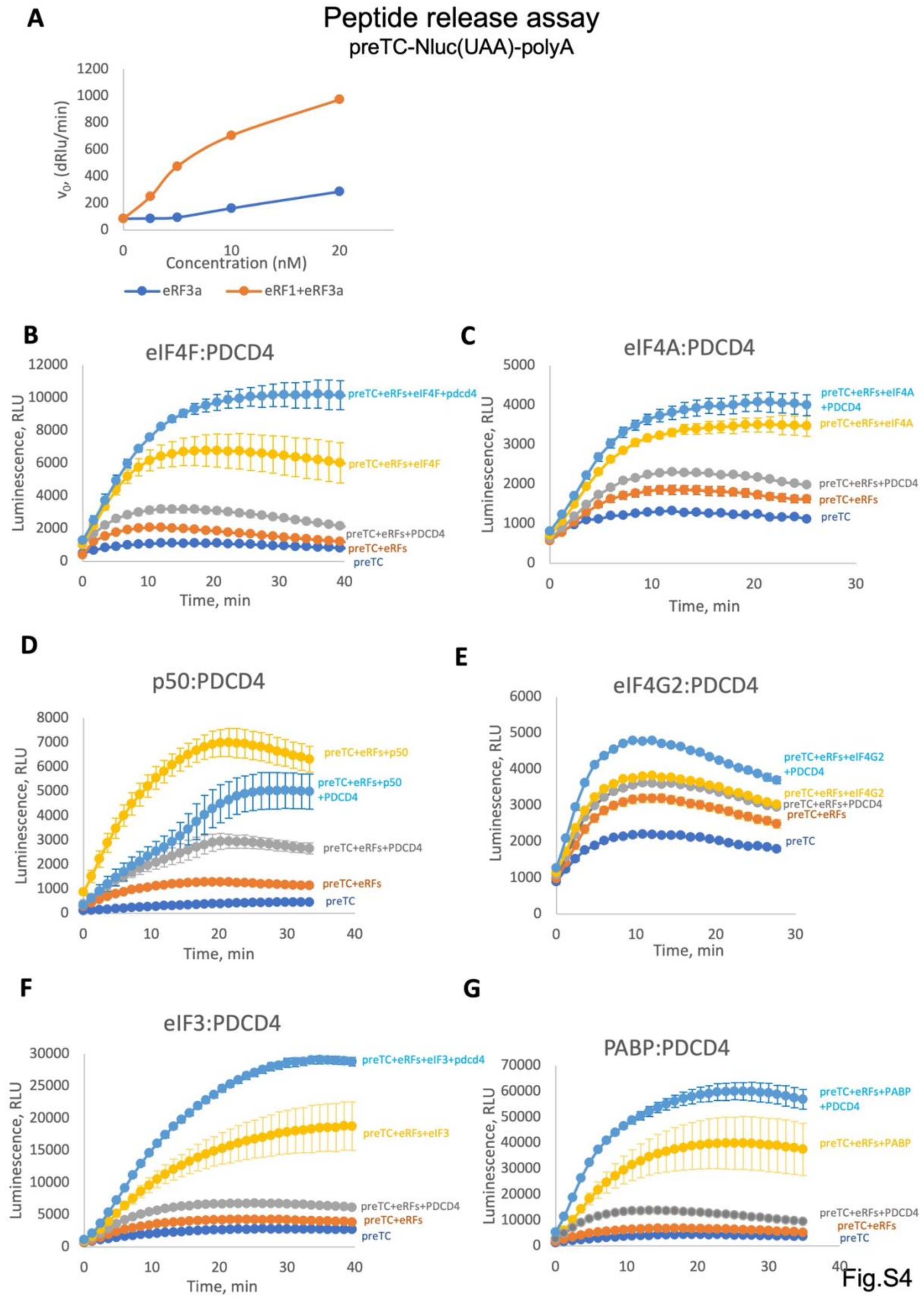

A
Peptide release assay
preTC-Nluc(UAA)-polyA
v0, (dRlu/min)
Concentration (nM)
eRF3a
eRF1+eRF3a
B
eIF4F:PDCD4
Luminescence, RLU
Time, min
preTC+eRFs+eIF4F+pdcd4
preTC+eRFs+eIF4F
preTC+eRFs+PDCD4
preTC+eRFs
preTC
C
eIF4A:PDCD4
preTC+eRFs+eIF4A +PDCD4
preTC+eRFs+eIF4A
preTC+eRFs+PDCD4
preTC+eRFs
preTC
D
p50:PDCD4
preTC+eRFs+p50
preTC+eRFs+p50 +PDCD4
preTC+eRFs+PDCD4
preTC+eRFs
preTC
E
eIF4G2:PDCD4
preTC+eRFs+eIF4G2 +PDCD4
preTC+eRFs+eIF4G2
preTC+eRFs+PDCD4
preTC+eRFs
preTC
F
eIF3:PDCD4
preTC+eRFs+eIF3+pdcd4
preTC+eRFs+eIF3
preTC+eRFs+PDCD4
preTC+eRFs
preTC
G
PABP:PDCD4
preTC+eRFs+PABP +PDCD4
preTC+eRFs+PABP
preTC+eRFs+PDCD4
preTC+eRFs
preTC


Fig.S4
